# mtDNAcombine: tools to combine sequences from multiple studies

**DOI:** 10.1101/2020.03.31.017806

**Authors:** Eleanor F. Miller, Andrea Manica

## Abstract

Today an unprecedented amount of genetic sequence data is stored in publicly available repositories. For decades now, mitochondrial DNA (mtDNA) has been the workhorse of genetic studies, and as a result, there is a large volume of mtDNA data available in these repositories for a wide range of species. Indeed, whilst whole genome sequencing is an exciting prospect for the future, for most non-model organisms’ classical markers such as mtDNA remain widely used. By compiling existing data from multiple original studies, it is possible to build powerful new datasets capable of exploring many questions in ecology, evolution and conservation biology. One key question that these data can help inform is what happened in a species’ demographic past. However, compiling data in this manner is not trivial, there are many complexities associated with data extraction, data quality and data handling. Here we present the mtDNAcombine package, a collection of tools developed to manage some of the major decisions associated with handling multi-study sequence data with a particular focus on preparing mtDNA data for Bayesian Skyline Plot demographic reconstructions.

## Introduction

Understanding a species’ demographic past can help inform many questions in ecology, evolution and conservation biology. Consequently, there is a lot of interest in methods that are able to infer how a population’s size may have changed through time. Traditional methods relied on insight from the fossil record [1–3]. However, although fossils are informative about many species, including our own, they remain a limited resource with coarse geographic and temporal resolution. In contrast, genetic methods have the potential to offer better resolution and are now established as the primary means by which a population’s past can be interrogated.

Mitochondrial DNA (mtDNA) has been used widely for demographic reconstruction. The haploid nature of mtDNA along with its rapid rate evolution [4], lack of recombination [5] and uniparental mode of inheritance [6] make it more sensitive to capture changes in population size than slower evolving nuclear genes [7] (Fig. 1). MtDNA therefore has the temporal resolution to capture the impacts of relatively recent events that might be of interest, such as the Last Glacial Maximum (LGM). In combination with coalescent-based reconstruction methods such as Bayesian Skyline Plots (BSPs) [8], mtDNA can be used to estimate a detailed population profile that stretches back tens, or even hundreds, of thousands of years. On the negative side, since the mtDNA genome does not recombine, it acts as a single locus and thus is subject to high levels of stochasticity, necessitating larger sample sizes of individuals than if multi-locus data were available.

**Figure 1.**
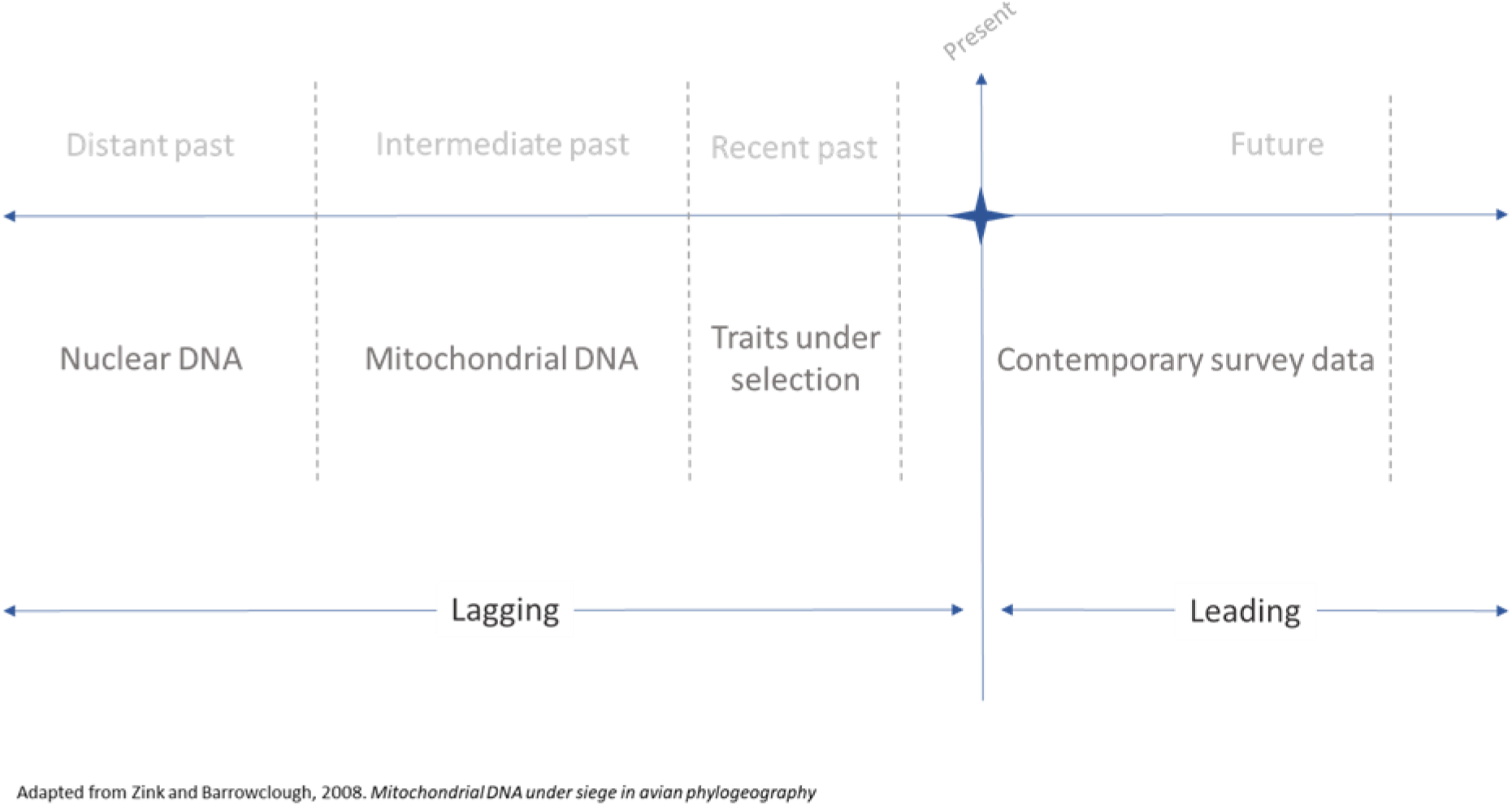
Utility of different loci for reconstructing different periods of population history.

With the falling costs of whole genome sequencing (WGS) and the growing interest in large scale sequencing projects, such as the Bird 10,000 Genomes Project (B10K) [9], the availability of WGS data is rapidly increasing. Using a single, high quality, diploid genome sequence, the pairwise sequentially Markovian coalescent (PSMC) method [10] can reconstruct a profile of population size through time for that species. However, PSMC is limited in its ability to capture details of population history more recently than ~1,000 generations ago [11]. The multiple sequential Markovian coalescent (MSMC), a method that builds on the PSMC framework, somewhat resolves this issue, using data from multiple individuals to improve the resolution of PSMC by an order of magnitude to more recent times [11]. However, this method is costly, requiring multiple, phased, high-quality genomes from the species of interest. Whilst phasing data may get easier as average sequenced read lengths increase, this is still a non-trivial step and phased data is frequently too difficult or costly to obtain for non-model species.

Whilst WGS is an exciting prospect for the future, for most non-model organisms’ classical markers such as mtDNA remain widely used [12]. Indeed, the falling costs of high throughput DNA sequencing, coupled with routine deposition of project data into public databases such as the National Centre for Biotechnology Information’s (NCBI) GenBank [13], has created a burgeoning resource of mtDNA sequence data. For the first time, these databases contain sufficient sequence data to allow users to build quality meta-datasets. Although individual studies may only be able to undertake spatially and temporally restricted sampling efforts, by creatively using pre-existing resources from multiple studies, it is now feasible to improve sampling strategy, range coverage and sample sizes without additional sampling. As the workhorse of population genetics studies for many decades, public domain mtDNA data are available in large numbers for a wide range of species across most higher taxa.

Although sequence databases are normally curated, data input is generally not standardised or error checked. Studies differ greatly in the length and identity of target sequence, the quality of sequence curation and, while some studies upload all sequences obtained, others merely upload unique haplotypes. There are also instances of incorrect sample assignation. Altogether, this means that to compile a comparable set of sequences from multiple studies requires extensive data processing. In the current paper, we consider the practicalities and problems faced by a meta-analysis of publicly available data and present the mtDNAcombine package. The mtDNAcombine package is a collection of tools developed to manage some of the major decisions associated with handling multi-study sequence data with a particular focus on preparing mtDNA data for BSP population demographic reconstructions (Fig. 2.).

**Figure 2.**
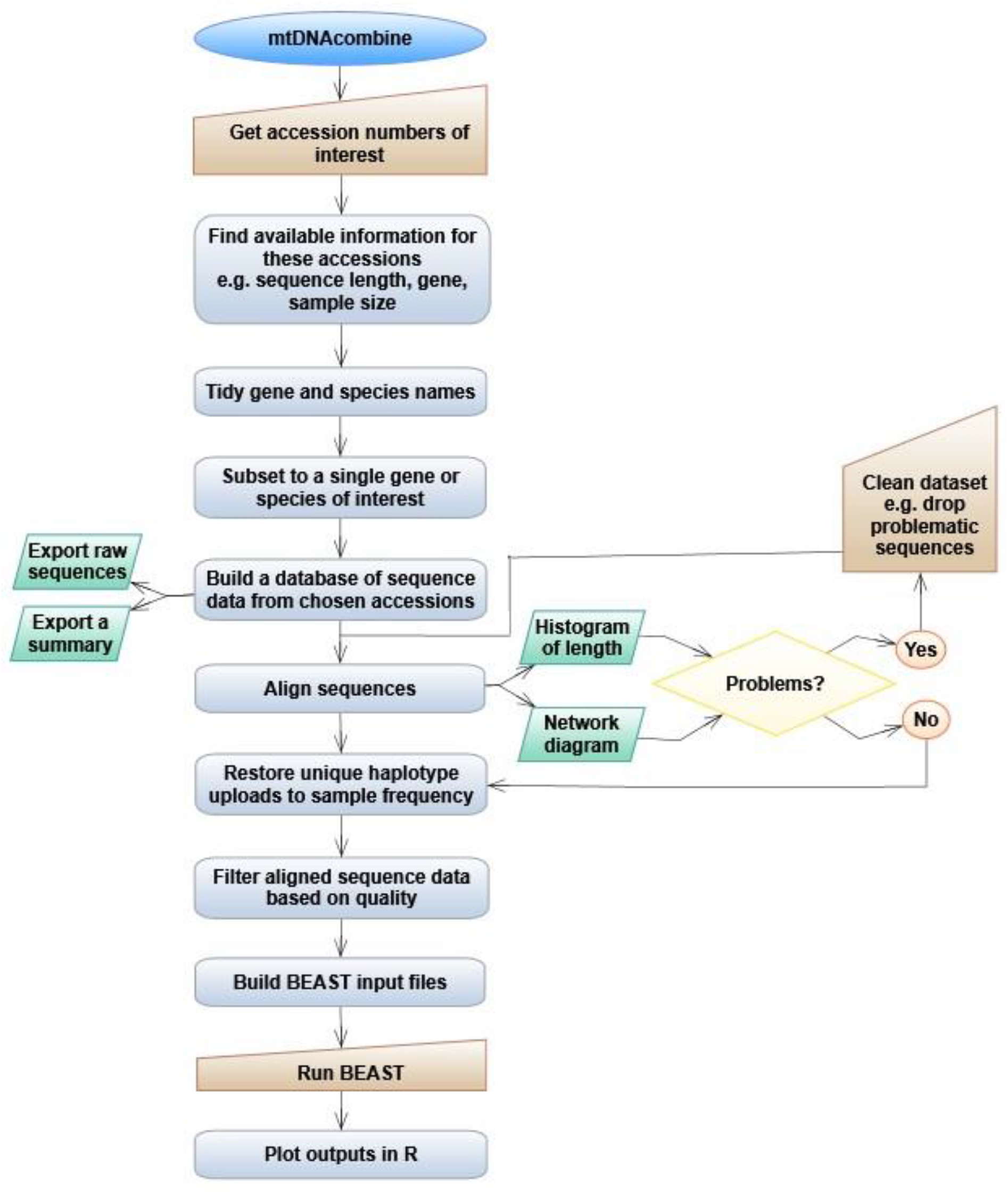
Flow diagram of mtDNAcombine pipeline showing decisions and steps supported by the package.

## Methods

### Data preparation

#### Raw data

Step one is to search annotated DNA databases to determine how many data sets are available. We focus on GenBank, which is the main public repository for mtDNA datasets. Their website is intuitive, and it is easy to set up a search for a given taxon. In mtDNAcombine, we import information (e.g. title of associated paper and sequence length) about relevant accessions into a dataframe with the ‘build_GB_dataframe’ function. We then proceed to explore and clean up this information to make it comparable across studies, and thus allow us to merge data for the any given species and create comparable datasets for multiple species.

It should be noted that, although GenBank staff review all submissions to GenBank, and quality control checks are performed before release, there is no standardised format for entering descriptive information. As a result, features such as alternative abbreviations for gene names, deprecated species names, subspecies names, and simple misspellings are all common. When nomenclature does not match between entries, filtering a large database for comparable samples is complex so, the mtDNAcombine pipeline includes two functions (‘standardise_gene_name’, ‘standardise_spp_name’) that allow the user to re-set common alternatives / errors in species and gene names to a chosen standard value.

#### Avoiding duplicate sequence entries

As BSP analysis draws information from haplotype frequency, it is important to try to avoid inclusion of duplicate entries because these can skew estimates of effective population size (*N_e_*) and alter the reconstructed timings of demographic events. Repeated entries for a single sample can come from multiple sources, for example, the NCBI Reference Sequence (RefSeq) project [14] aims to curate records and associated data, providing a set of reference standards. As these data are drawn from the International Nucleotide Sequence Database Collaboration (INSDC, which consists of GenBank, the European Nucleotide Archive (ENA), and the DNA Data Bank of Japan (DDBJ)) databases, a basic search can recover two accessions for the same sample; the RefSeq accession and the source record(s). In this instance, the duplicates can be distinguished because all RefSeq records include an underscore (“_”) in their accession number, while simple repository accessions never have this character. Our code (‘load_accession_list’ function, called within ‘build_GB_dataframe’) will automatically (and silently) remove any RefSeq record if the original accession is also found to be present in the dataset; however, users should be aware that these exclusions are being made.

Duplications can also arise from re-uploaded / re-sequenced samples. This occurs most frequently when multiple studies sample a single museum specimen, though there are other scenarios which can lead to a single individual being sequenced by multiple studies. Re-sequenced samples are often hard to identify and recognising repeated use of published alternative ID numbers (such as specimen numbers) are sometimes the only indications that the same individual has been sequenced by multiple studies. Although an occasional duplicate entry in a moderate sample size of around 100 sequences is unlikely to cause a significant skew in the recovered population history, authors should be conscious that this source of duplicate entry exists and needs to be avoided whenever possible. Unfortunately, there is no simple programmatic way to avoid it given the information provided in GenBank.

#### Alignment

After sequence data have been obtained, they must be aligned. A number of public domain software programs are available that can achieve this, including T-Coffee [15], MUSCLE [16,17] and MAFFT [18]. In mtDNAcombine, we chose to use ClustalW [19], implemented through the R package ‘msa’. [20]. Though BEAST can handle missing / ambiguous bases [21], we consider it best to use alignments without gaps or ambiguities. Whilst some insertions or deletions may be genuine, when working with sequences from multiple sources, the data are likely to have been sequenced with different techniques to varying standards. Inclusion of basic sequencing errors could drive miscalculations in later analyses and the volume or type of errors will not be consistent across all studies, nor across all taxa. We therefore recommend that, to ensure consistent sequence quality, all sites with ambiguities, insertions, deletions and missing data should be removed. This is done automatically within the ‘align_and_summarise’ function in mtDNAcombine.

#### Diagnostic plots

Compiling data from multiple studies produces a series of known challenges which we tackle individually in the following sections. The ‘align_and_summarise’ function draws a series of key diagnostic plots for each species dataset being handled. These plots are designed to help the user quickly visualise the data, enabling rapid identification of any problems in the aligned data. If these diagnostic plots look problematic, it is then possible to return to the original input files and revaluate the raw sequence data on a case-by-case basis. The user can then decide to proceed with the analysis, return to the pipeline with an edited set of samples, or choose to drop the dataset entirely if too many samples / studies have to be excluded.

#### Sequence length

For any group of studies there will be numerous reasons the samples were original collected and sequenced. Each project will have had, among other things, a different budget, time constraints, target area of the mitochondrial genome, and available sequencing technology, meaning that different lengths of the genome / target gene will have been sequenced. In some instances, only very short sections of the gene of interest will have been sequenced. If the number of base pairs (bp) is too low, the sample is unlikely to hold enough information to be informative for population demographic reconstruction. The ‘align_and_summarise’ function will drop individual accessions that are below a user-set threshold before processing the data. There can be no out-of-the-box value for this ‘minimal length’ as the most appropriate size will vary with a wide range of factors such as the gene under investigation, mutation rate, absolute gene length, and the available sample size. However, excluding any samples that clearly hold insufficient information before aligning and cropping sequences to the maximum overlapping area prevents an excessive loss of information if one very short sequence were included.

Equally, above the minimal length that has been set, there can still be a wide variance in the number of base pairs, or region of the focal gene sequenced by different studies. Automatically cropping all the sequences to the maximum overlap length may result in the loss of a large amount of data unbeknownst to the user. Therefore, in order that the process of alignment and sequence trimming is transparent, one of the diagnostic plots mtDNAcombine produces is a histogram showing the original variation in sequence length as well as the length of the trimmed, maximum overlap, dataset (Fig. 3, vignette section ‘Diagnostic plots’). This plot flags instances where a large number of base pairs have been removed in order to include a shorter sequence. Sequence length versus sample size is a trade-off that individual users may want to weight differently depending on the data available. By presenting the information, mtDNAcombine allows the user to go back, review, and revise the input data if they want.

**Figure 3.**
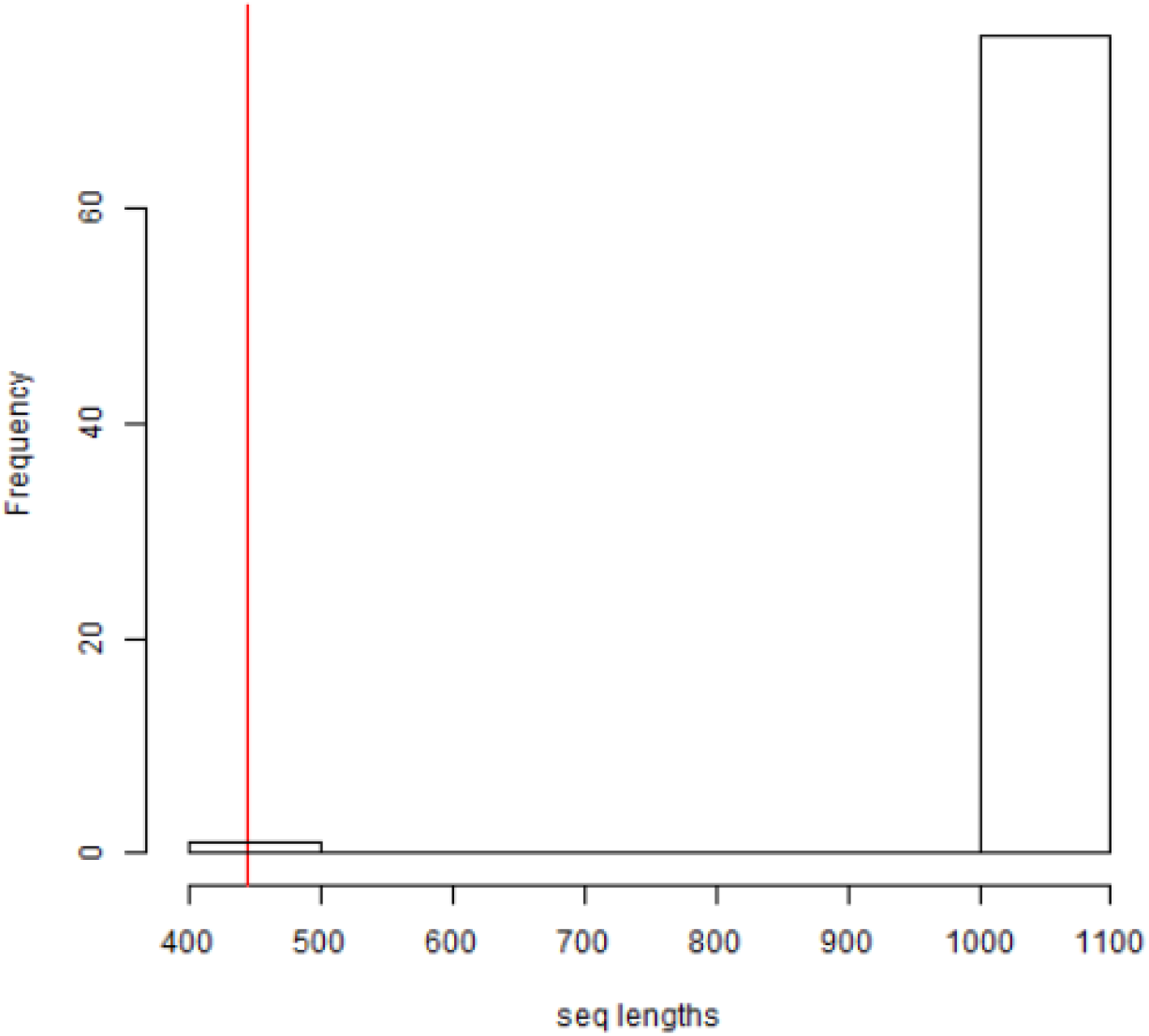
Example of diagnostic plot for sequence trimming in the ‘align_and_summarise’ function. Histogram shows that, in order to trim all sequences to the maximum overlapping length (red line), the majority of samples have had to be heavily cropped.

#### Haplotype frequency

Studies differ in the ways they deposit data. Some upload a single copy of each haplotype they found, while others upload sequences for each individual sampled. Datasets built exclusively of unique haplotypes are not suitable for a BSP analysis [22]. Where only unique haplotypes have been uploaded, it is vital to find the number of samples these haplotypes represent, or the study must be excluded. Routinely checking every source publication to see whether they uploaded only a single copy would be tedious and may become impractical for larger analyses. To guide this process, the ‘align_and_summarise’ function flags studies in which all haplotypes are unique (i.e. there are no replicates) as candidates for further investigation. A text file of individual accession numbers is also produced, including a column for the user to input new frequency information. Once satisfied that the sampled frequency for each haplotype has been recorded correctly within this document, the table can be read back into R, and the function ‘magnify_to_sampled_freq’ will build the dataset up to correct sample sizes. See vignette section ‘Haplotype frequency’ for a worked example.

#### Population Structure

Population sub-structure is known to cause problems for demographic reconstructions methods and BSP analysis is no exception [23–26]. BSP analysis, like other coalescent methods, is founded on the Wright-Fisher model and hence assumes panmixia [27]. This assumption is violated by population sub-structure [23,28], which acts to reduce the probability that lineages from different demes coalesce. In practice, depending on the sampling strategy employed, sub-structure can lead to inflated population size estimate in older parts of the reconstructed history but can also noticeably reduce apparent population size at the present [23]. Accurate demographic reconstruction therefore requires careful consideration of whether sub-structure is or might be present.

Once DNA sequences have been identified, downloaded, aligned, and multiplied up to sampled frequency, the level of population structure can be assessed. One of the most intuitive approaches is to visualise the haplotype network diagram for each dataset. To maintain a streamlined approach, we draw network diagrams within R using the package ‘pegas’ [29]. These network diagrams are one of the diagnostic plots created by the ‘align_and_summarise’ function (vignette section ‘Network diagram’).

Depending on the level of supplementary detail available for each sample, the decision to split a population for analysis can be simple. For example, in instances where sampling location data are available and clear geographic divisions coincide with major genetic clades, datasets can be separated and multiple sequence files handled as individual datasets. However, it is important not to over-split the data. Clades are a natural feature even of fully homogeneous populations, so if any obvious clades are removed, what is left will tend to be star-like haplotype clusters. Such clusters will often yield a signal of population expansion which may or may not be real. Deciding if and where to divide datasets remains one of the more subjective and difficult challenges and it can be worth investing time into running data sub-sets to determine the impact of alternative splitting decisions.

#### Outliers

We frequently found instances of extreme outliers, single haplotypes that were separated from all others by many base changes. Such outliers may be genuine but equally may reflect immigrant individuals, sample mislabelling [30], amplification of integrated nuclear copies, incorrect accession codes, or even result from poor-quality sequencing. We feel that the benefits of including these outliers in case they are genuine are far outweighed by the risk that they distort the process of inference. We therefore recommend that outliers are identified and removed, although it is useful to retain copies of the original files so that the impact on inferred demographic histories can later be investigated if necessary. Within the ‘outliers_dropped’ function, any “extreme outliers” are removed from the working dataset. We recognise that factors such as species life history, species population history, data availability, and data quality will influence the criteria for data inclusion. Therefore, the degree of separation from other haplotypes necessary for a sample to be classified as an “extreme outlier” is something that can be set by the user.

### Setting up and running BEAST

#### BEAST input

In large comparative studies, as many steps as possible should be kept constant. This minimises the chance that the analysis becomes prohibitively time-consuming and helps to make the outputs as directly comparable as possible. The process of setting up and parameterising a BSP analysis in BEAST is well-described in several papers as well as in the accompanying textbook [21] so we will not go into detail here. Briefly, BEAST requires values for a range of parameters of which arguably the most important is mutation rate. Selection of an appropriate mutation rate is a persistent problem in genetic studies. With BSP analyses, mutation rate influences the scaling of both inferred population size and timing of events, but it does not affect the overall profile shape. Both the mutation rate itself and its associated confidence will vary between taxa and it is necessary for the user to consider how best to standardise this to maximise consistency across profiles. For certain groups, attempts have been made to provide rates for a large number of taxa [31], though this kind of resource is far from universal as yet.

To maximise the probability that a given run converges, it can be a good idea to use fairly tight constraints on initialising parameters such as the number of population size changes. This decision will be study-specific with no one-size-fits-all approach. Moreover, changing priors and parameter values can alter outputs and should be done in accordance with best Bayesian practices [21]. Bearing this in mind, we suggest that a loss of resolution in some profiles may be a necessary trade-off if the maximum number of species is to be included.

The mtDNAcombine package function ‘setup_basic_xml’ utilises the ‘babette’ package [32] to build basic XML files form the data set processed earlier in the pipeline. The skeleton XML files will need editing (e.g. defining mutation rate, model choice, output names) but their creation minimises the number of steps the user needs to perform manually, speeding up the process and reducing the opportunity for the introduction of human error. Once parameterisation decisions have been made and the XML input files finalised, whenever possible, we encourage use of the BEAGLE library [33] when running BEAST2, since this can significantly improve the speed of a run.

#### BEAST output

Interpretation of BEAST outputs has been covered well in the literature e.g. [22,23] and by those who designed and built the software [34–38]. As with any statistical model, checks need to be done to confirm the reliability of the output. In BEAST2 these are generally undertaken using the software package Tracer [39] and focus on appropriate convergence of the Markov chain.

As a rule of thumb, outputs should be treated with caution wherever the effective sample sizes (ESS) for a given parameter drops below 200. Similarly, duplicate runs should be used to confirm that the posterior probability distributions stabilise at similar values. Whilst ESS values can be captured directly through the package ‘babette’ [32], we think that a visual inspection of each run in Tracer is best practice. Whilst doing so, it is then possible to export extensive summary data from the ‘Bayesian Skyline Reconstruction’ tab (found under ‘Analysis’ in Tracer). These Tracer exports are detailed, informative, and concise to work from, ideal for tasks such as downstream data visualisation as we do in mtDNAcombine.

#### Plotting profiles in R

BSPs can be drawn using the programme Tracer [39]. However, for more flexibility, and to facilitate exploration of the profiles in greater detail, we chose to visualise the reconstructed profiles in R. Within the mtDNAcombine package vignette, we present example code for plotting Tracer output data as BSP profiles (section ‘Exploring outputs’). However, it is anticipated that data presentation will be highly project specific, therefore this code is not tied up in functions, enabling easy editing and adaptation by the user.

#### Cautions

Skyline plots offer a powerful tool set but are easily over-interpreted. Although covered in several recent reviews [22,40], over-interpretation continues to be an issue and hence its dangers are worth re-iterating. Unsurprisingly, problems are greatest with weaker data: smaller sample sizes, uneven sampling strategy, and / or when drawn from a species with strong population substructure [22,23]. For example, an investigation of the same species, the common rosefinch, based on two mtDNA datasets with very different sample sizes gives us contrasting results (Fig. 4.). The smaller sample set, cytb, suggest a weak linear increase in size over time but the larger dataset, ND2, uncover a rapid, almost 100-fold increase in size. This clearly indicates that interpretation of BSP plots must be done with appropriate consideration for the data quality.

**Figure 4.**
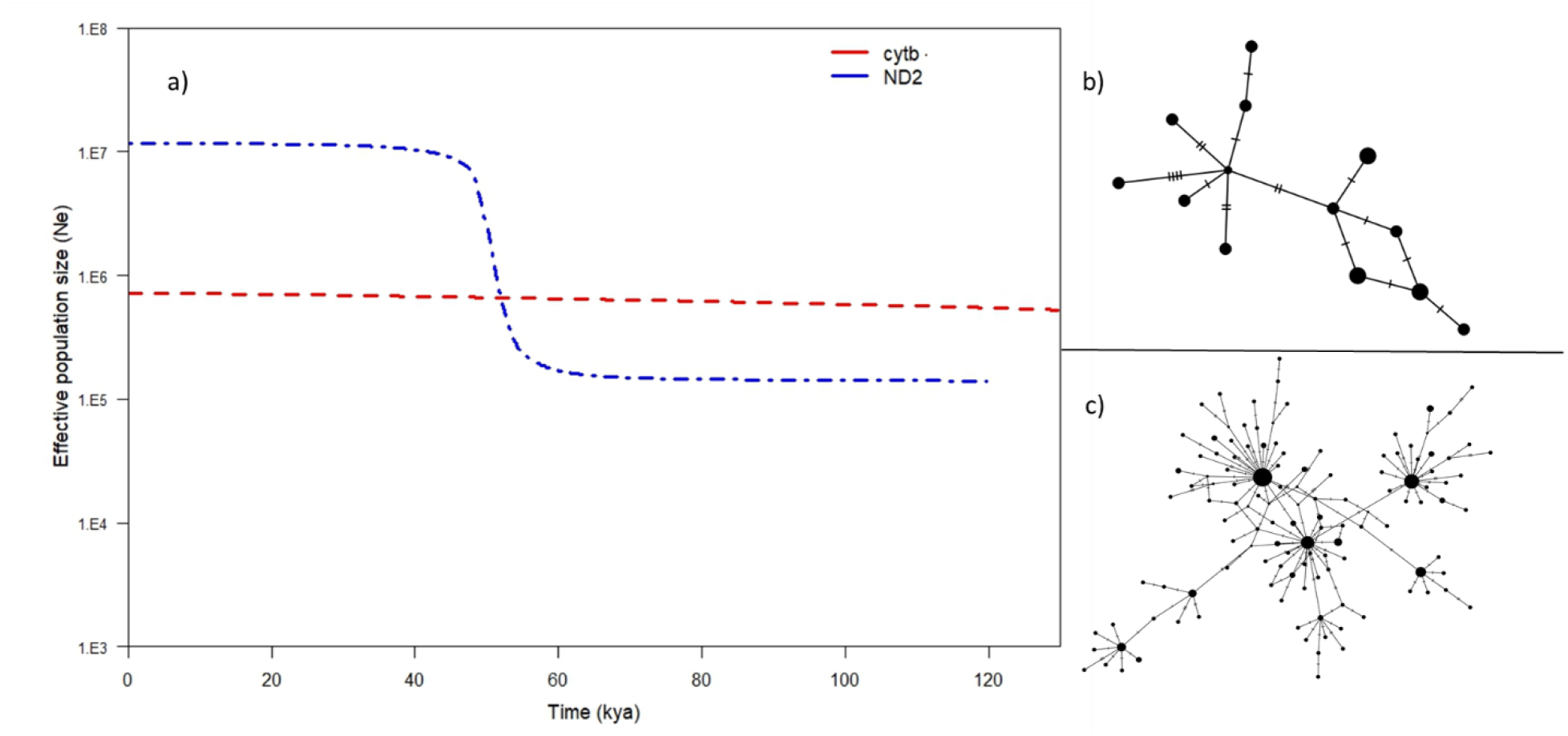
Comparison of two dissimilar BSP profiles drawn from different mtDNA datasets of the common rosefinch. a) Red line is median value for cytb BSP profile, blue line is median value for ND2 BSP profile. The cytb dataset includes 15 samples, ND2 dataset 190 samples. The varying levels of information available for inferences to be drawn from are clearly shown in b) the median joining network (MJN) for cytb dataset, and c) MJN for ND2 dataset.

#### Uploading sequence data

When assembling large annotated DNA databases using published data, many sequences are ‘lost’ due to inaccuracies or inconsistencies in how the data are uploaded to repositories. Unless the accession process becomes more standardised, idiosyncrasies and errors will continue to render an appreciable proportion of the potential data unusable. We therefore encourage people who wish to upload data to take the time to complete as many supplementary fields as possible and to be sure they undertake basic formatting checks such spell-checks, correct capitalisation and use of standard abbreviations. Where accompanying information is not uploaded to repositories, we urge authors to make this information easily accessible to readers. For example, downstream use will be facilitated by providing haplotype frequency data or detailed sampling location data as supplementary files (ideally well formatted text files which are easy to process) rather than embedded tables or images within manuscripts.

## Conclusions

With the exponentially expanding volume of data in public DNA sequence repositories, there is now more genetic information available than ever before. Building large meta-data sets by combining existing data offers the opportunity to explore new and exciting avenues of research e.g. [41–43]. However, compiling multi-study datasets still remains a technically challenging prospect. Unknown sequence quality, little to no control over sampling structure, potential errors in species identification, and limited control of sample size are all factors that can negatively affect a comparative study if not carefully handled.

Here we present the mtDNAcombine package, providing a pipeline to streamline the process of downloading, curating and analysing mitochondrial sequence data (Fig. 2). At the moment, the lack of standardisation in the data upload process exacerbates the inevitable complexities of combining data from multiple origins. Whilst some samples, sequenced early in the molecular era, are allowably poorly documented we urge people to be careful when uploading data today. The more information about a sample that is included online, alongside sequence data, the more likely that sequence will be usable by others. Equally, with the volume of data available today the accuracy of associated meta-data and sequence tags / labels is vital for ensuring the data are retrievable when broad, automated, searches are used. We suggest that a focus on quality control for additional information about each sample will make a noticeable difference to the ease with which public databases can be mined for relevant information and this exceptional resource exploited. We hope that our discussion, whilst highlighting common pitfalls, provides solutions and suggestions to guide the process of compiling data sets from online databases.

## Data Availability

mtDNAcombine source code and full vignette can be found at: https://github.com/EvolEcolGroup/mtDNAcombine. The data used as examples in this paper were derived from publicly available data in GenBank (https://www.ncbi.nlm.nih.gov/genbank/). A sample dataset is available within the R package.

## Author contributions

E.F.M. performed the analyses, prepared all figures and wrote the manuscript. All authors conceived the idea and design, as well as reviewing the manuscript.

## Additional information

The authors declare no competing financial interests.

